# Markov chain models of cancer metastasis

**DOI:** 10.1101/263350

**Authors:** Jeremy Mason, Paul K. Newton

## Abstract

We describe the use of Markov chain models for the purpose of quantitative forecasting of metastatic cancer progression. Each site (node) in the Markov network (directed graph) is an organ site where a secondary tumor could develop with some probability. The Markov matrix is an N x N matrix where each entry represents a transition probability of the disease progressing from one site to another during the course of the disease. The initial state-vector has a 1 at the position corresponding to the primary tumor, and 0s elsewhere (no initial metastases). The spread of the disease to other sites (metastases) is modeled as a directed random walk on the Markov network, moving from site to site with the estimated transition probabilities obtained from longitudinal data. The stochastic model produces probabilistic predictions of the likelihood of each metastatic pathway and corresponding time sequences obtained from computer Monte Carlo simulations. The main challenge is to empirically estimate the N^2 transition probabilities in the Markov matrix using appropriate longitudinal data.

## I. Introduction to the Type of Problem in Cancer

Predictive mathematical models of cancer progression for the purposes of quantitative forecasting rely heavily on the ability to obtain appropriate longitudinal data of cohorts of patients with different tumor types whose disease progresses over time (say 5-20 years depending on the cancer type) undergoing different treatment modalities all of whom start out with non-metastatic disease. These data are then used to determine parameters (transition probabilities in a Markov matrix) in a dynamical (typically stochastic) progression model that can then be used (i) to make forward quantitative predictions and to quantify the uncertainty of the predictions; (ii) develop Monte Carlo simulations to create distributions of computer generated patients with correct statistical properties; (iii) run computational clinical trials to test hypotheses and pin down causality. The relevant mathematical modeling techniques are much further developed in financial prediction settings [1] and in weather forecasting modeling [2, 3] but lag considerably farther behind in disease forecasting applications, partly because of insufficient and low-quality data (by comparison) and partly because the relevant biological mechanisms are not as well understood [4]. One area where recent progress has been made is in the development of Markov chain predictive models [5-9] of cancer metastasis, where the underlying driver of the dynamics is an N x N transition matrix made up of N^^^2 transition probabilities which serve as the main parameters that must be estimated [10, 11] with appropriate data.

Figure 1 is a spatiotemporal progression diagram which organizes a longitudinal data set (as described in [9]) in a form useful for the estimation of the various transition probabilities that populate a Markov matrix. The inner most ring represents the primary tumor (breast), the second ring out shows the distribution of first metastatic sites, with sector sizes corresponding to the percentage of patients with first metastases at each of the given sites. Each of the subsequent concentric rings represents the distribution of additional metastatic tumors. The black sector at the end of a given ray indicates that the patient is deceased. Following along a given ray from the center of the diagram lays out a particular metastatic pathway (chronological sequence of metastatic tumors) for a given patient. By computing the probabilities of transitioning from site-to-site in these spatiotemporal diagrams, we can estimate the transition probability of the cancer spreading from any given site to another in one step [9]. This information gives rise to a Markov transition matrix which forms the dynamic driver of the model. Markov models have been used extensively in other medical settings both for survival estimation [12, 13], as well as for tumor progression [14-19] and more general applications [20]. Network models of disease progression and spread have also been developed in other contexts [21, 22].

**Figure 1.**
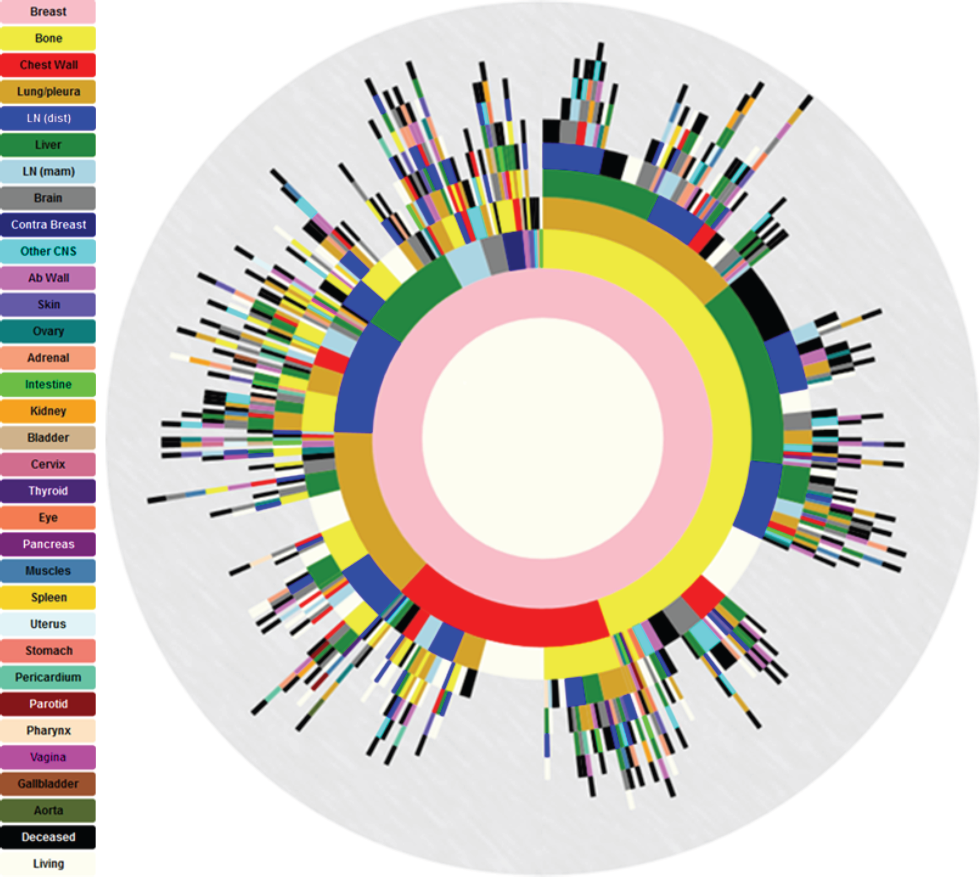
Spatiotemporal progression diagram of 446 primary breast cancer patients [9]. The innermost to outermost rings show progression patterns from primary breast (pink ring) to distant metastatic sites (subsequent rings). Circular arc length of each sector represents the percentage of patients with a metastatic tumor in that location.

## II. Illustrative Results of Application of Methods

Once the transition probabilities from site to site are obtained via appropriate interpretation of the data in Figure 1 (the main approximation is the Markov assumption of using all data in the diagram associated with patients that progress from site A to site B regardless of the ring number), the probabilities of each of the patients’ metastatic pathways can be computed by multiplying the appropriate sequence of transition probabilities. A common pathway for metastatic breast cancer, for example, is breast ➜ bone ➜ liver ➜ deceased, as represented by the pink inner ring, followed by the yellow (bone) second ring sector, then the green (liver) third ring sector, and finally by the black (deceased) sector. Calculating the probabilities of each of the sequences present in the data set allows us to rank order each pathway by likelihood, and thereby make a cutoff as to how many of the most likely pathways to use in the model. Figure 2, for example, shows a reduced Markov diagram associated with the top 30 two-step pathways, aggregating all of the breast cancer subtypes and treatment modalities into one group. Each site can be categorized as a potential spreader site (red) or a sponge site (blue) based on the ratios of the probabilities of the paths out compared to the paths in [6, 9]. For metastatic breast cancer, bone is the predominant spreader site, while lung/pleura is the predominant sponge site. We can further subdivide the data into the four common breast cancer sub-types, which largely determines treatment modality and survival: ER+/HER2+; ER+/HER2-; ER-/HER2+; ER-/HER2-. Figure 3 depicts the reduced Markov diagrams associated with each of these sub-types.

**Figure 2.**
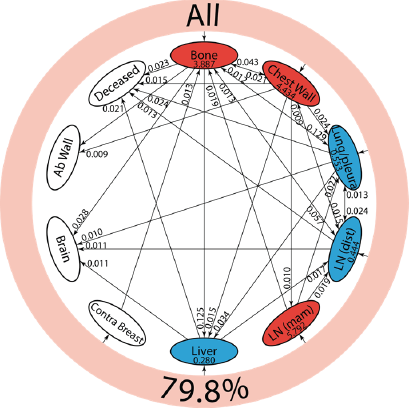
Reduced Markov models showing the top 30 two-step pathways emanating from primary breast (pink ring) [9]. Pathway probabilities are shown at the end of the second step, designated by an arrow pointing into a node. Nodes are classified as “spreader” (red) or “sponge” (blue) based on the ratio of their cumulative incoming and outgoing two-step probabilities. Spreader and sponge factors are listed inside each respective node’s oval.

**Figure 3.**
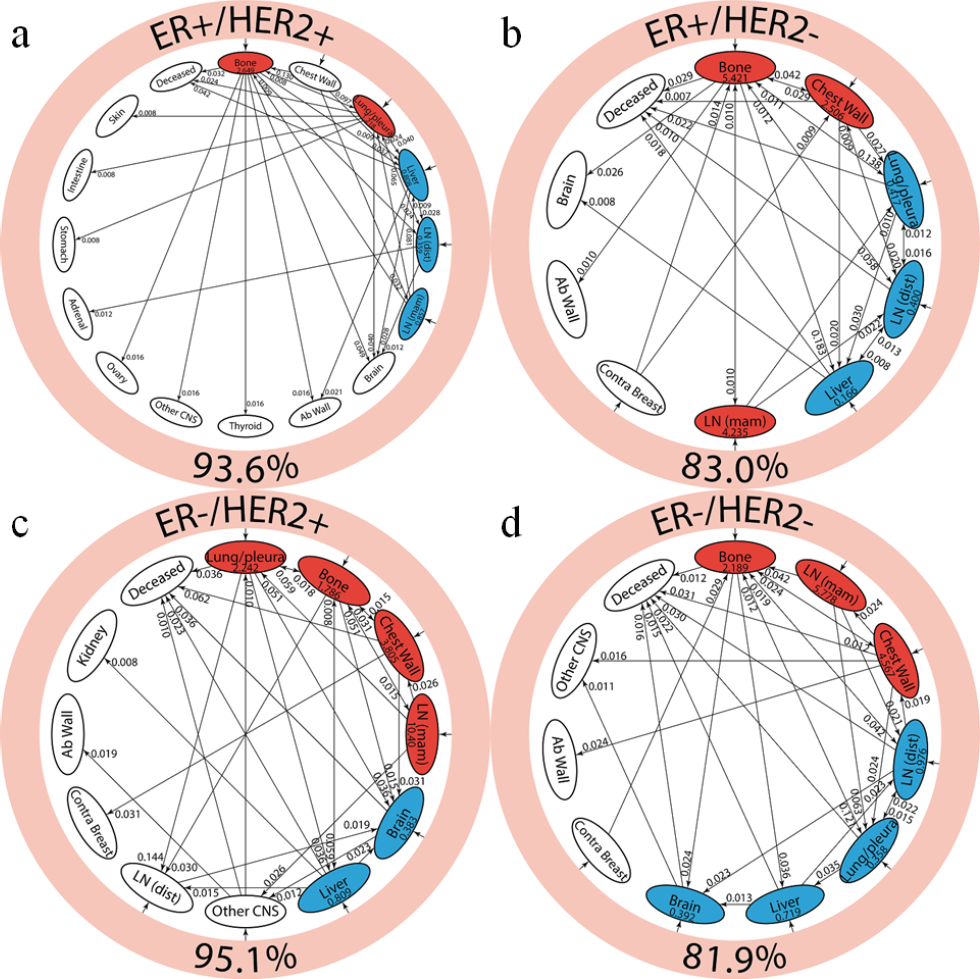
Reduced Markov models showing the top 30 two-step pathways of hormonal subgroups of primary breast cancer [9]. Lower number indicates the % that the 30 pathways capture. (a) ER+/HER2+ breast cancer, (b) ER+/HER2-, (c) ER-/HER2+, and (d) ER-/HER2-.

Markov models of complex dynamical processes, despite their step-to-step simplified assumptions (i.e. no history dependence), retain their appeal as a first approach to modeling spatiotemporal dynamics because of their ease of interpretability, the clarity of the resulting dynamics, and their use in isolating phenomena on which to invest more effort into building more elaborate models involving nonlinear systems of ordinary differential equations, partial differential equations, or hybrid systems.

## III. Quick Guide to the Methods (1 Page)

A discrete Markov chain dynamical system is governed by the equation:

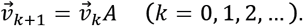

*A* is an N x N transition matrix comprised of transition probabilities, *P_ij_*, that give the probability of going from state *i* to state *j* at each step. The matrix is row stochastic:

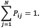

The state vector, 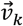, contains the probabilities of metastatic tumors developing at specific locations (summing to 1) at a given time step k. An initial state vector, 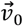 (k=0) is represented with a 1 in the position for the primary tumor location and 0’s elsewhere. Then:

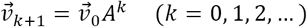

indicating that the underlying dynamics that defines disease progression is interpreted as a weighted, random walk on directed graph defined from the transition matrix. A more detailed description can be found in [11,20].

